# Direct Metatranscriptome RNA-seq and Multiplex RT-PCR Amplicon Sequencing on Nanopore MinION – Promising Strategies for Multiplex Identification of Viable Pathogens in Food

**DOI:** 10.1101/700674

**Authors:** Manyun Yang, Alyssa Cousineau, Xiaobo Liu, Daniel Sun, Shaohua Li, Tingting Gu, Luo Sun, Yaguang Luo, Mingqun Xu, Boce Zhang

## Abstract

Viable pathogenic bacteria are major biohazards that pose a significant threat to food safety. Despite the recent developments in detection platforms, multiplex identification of viable pathogens in food remains a major challenge. A novel strategy is developed through direct metatranscriptome RNA-seq and multiplex RT-PCR amplicon sequencing on Nanopore MinION to achieve real-time multiplex identification of viable pathogen in food. Specifically, this study reports an optimized universal Nanopore sample extraction and library preparation protocol applicable to both Gram-positive and Gram-negative pathogenic bacteria, demonstrated using a cocktail culture of *E. coli* O157:H7, *Salmonella enteritidis*, and *Listeria monocytogenes*, which were selected based on their impact on economic loss or prevalence in recent outbreaks. Further evaluation and validation confirmed the accuracy of direct metatranscriptome RNA-seq and multiplex RT-PCR amplicon sequencing using Sanger sequencing and selective media. The study also included a comparison of different bioinformatic pipelines for metatranscriptomic and amplicon genomic analysis. MEGAN without rRNA mapping showed the highest accuracy of multiplex identification using the metatranscriptomic data. EPI2ME also demonstrated high accuracy using multiplex RT-PCR amplicon sequencing. In addition, a systemic comparison was drawn between Nanopore sequencing of the direct metatranscriptome RNA-seq and RT-PCR amplicons. Both methods are comparable in accuracy and time. Nanopore sequencing of RT-PCR amplicon has higher sensitivity, but Nanopore metatranscriptome sequencing excels in read length and dealing with complex microbiome and non-bacterial transcriptome backgrounds. To the best of our knowledge, this is the first report of metatranscriptome sequencing of cocktail microbial RNAs on the emerging Nanopore platform. Direct RNA-seq and RT-PCR amplicons sequencing of metatranscriptome enable the direct identification of nucleotide analogs in RNAs, which is highly informative for determining microbial identities while detecting ecologically relevant processes. The information pertained in this study could be important for future revelatory research, including predicting antibiotic resistance, elucidating host-pathogen interaction, prognosing disease progression, and investigating microbial ecology, etc.

## Introduction

Biological threats, including bacteria, viruses, and parasites, remain as the top food safety challenge in the United States. According to the CDC surveillance for foodborne disease outbreaks most recent annual report, in 2016 outbreaks attributed to bacterial infection comprised 44% of the total 645 outbreaks and caused 76% of the 847 hospitalization cases (1). More importantly, a recent report published by U.S. Department of Agriculture Economic Research Service (USDA ERS) stated that food safety challenges caused an annual loss of $15.5 billion to the economy and the top 10 infectious bacteria alone contribute to $10 billion in economic loss (2). These statistics revealed that bacterial infection is the primary concern among all biological threats. To cope with the threat of bacterial infection to public health, the demand for a rapid and highly sensitive method to detect and identify bacterial pathogens in food is enormous and becoming more urgent, especially after the implementation of the Food Safety Modernization Act (FSMA) in 2011.Commercial food safety testing methods include traditional plate counting methods, immunological techniques such as enzyme-linked immunosorbent assay (ELISA), lateral flow immunoassay chip, and electrochemical biosensors, chromatography, as well as nucleic acid-based approaches (3). As summarized in Table S1, the widely recognized cultivation methods and commercial rapid detection systems that are available for food defense applications have major limitations, such as large sample size, long turnaround time, and intensive labor demands. Rapid detection systems also failed to address unique changes to food safety and food defense, despite the recent success in medical and clinical diagnostics.

There are two major unique challenges in food defense, especially in identifying biohazards in food. First, viable bacteria are the etiological agents of foodborne illnesses. Most rapid detection methods have limited discretionary power to identify bacterial viability (summarized in Table S1). Current genome-based technology includes polymerase chain reaction (PCR), real-time PCR (qPCR), fluorescence *in situ* hybridization (FISH), nucleic acid sequence-based amplification (NASBA), and loop-mediated isothermal amplification (LAMP). PCR based methods are sensitive and specific but easily generate cross contamination between pre-PCR and post-PCR products. Insufficient permeability of cell walls and the inherent autofluorescence of the substrate will decrease the efficiency of FISH (4). NASBA and LAMP do not require a thermocycler, however, NASBA shows a size range limitation of target RNA and LAMP requires complex primer design that cannot be adapted to multiplex amplification (5). In addition, all these approaches suffer from the limited capability to identify viable pathogens (6). False positive results are a major issue for DNA-based approaches. This is due to the inability to differentiate DNA molecules in viable bacterial cells from the genomic background, which is comprised of stable DNA molecules from the microbiota, the food matrices, and dead pathogens inactivated during food processing and storage (7, 8). In contrast, transcriptome-based technologies which utilize RNA as alternative biomarkers for bacterial viability hold more promise, because RNA molecules tend to have a shorter half-life than DNA in the environment when cells are inactivated (7, 8). Recent progress was made using reverse transcription PCR (RT-PCR), but false positives also plague RT-PCR approaches (9). This can be explained by non-specific amplification of RNA molecules from food matrices and microbiota (10, 11). Subsequent sequencing of the RT-PCR amplicons has the potential to significantly improve the accuracy of the transcriptome-based approach by identifying the origin of the amplicons. The second prominent challenge is multiplex identification without the need for assay customization to each individual microbial threat. Each food commodity often faces multiple, and sometimes random, threats from dozens of major etiological agents (12). A monitoring and inspection system should entail capacities of multiplex identification. Nonetheless, conventional systems depend on the customization of recognition elements, like antibodies or enzymes, to achieve multiplex detection, which can be self-prohibitory economically (13, 14). Therefore, a feasible strategy should enable multiplex identification without the need to customize for individual threats, which can be of great importance and benefit to food defense. Several multiplex RT-PCR methods were developed for *S. aureus*, *Salmonella* and *Listeria* using food models over the last decade (15–20). However, a recent validation study suggests that multiplex RT-PCR may also generate false positive results in real food samples, especially if rRNA is the target template (11). Very recently, Next Generation Sequencing (NGS) platforms, such as Illumina, have emerged as a new strategy for food defense (21–25) but,its applications in food testing are very limited. NGS does not permit timely analysis, as these platforms generate sequence reads in parallel and not in series, so data analysis can add significant burden to total turnaround time. Additionally, NGS relies on non-portable and expensive equipment, which is also economically self-prohibitory for the food industry. The novel Oxford Nanopore MinION sequencer has emerged as a promising method of food pathogen detection based on its rapid, cost effective, portable, and high-throughput RNA and DNA sequencing workflows (26–29). Nanopore sequencing is a third-generation sequencing platform that can produce long reads on DNA and RNA molecules and perform real-time metagenomic and metatranscriptomic sequence analysis on the pocket-sized Nanopore MinION device (30). This technology can be used to identify viral pathogens, as well as microorganisms such bacteria and fungi (31–33). A few studies have demonstrated Nanopore’s potential for food safety application using metagenomic sequencing in clinical and food samples (34), however, like other genomic approaches, stable DNA molecules can cause false-positive identification, but the studies did not include a validation of whether the nanopore metagenomic sequencing data only correlates with viable pathogens. Direct RNA sequencing on Nanopore was successfully developed in 2018 on the Nanopore MinION (30).

Therefore, for the first time, RNA-enabled Nanopore sequencing is evaluated for its potential in achieving multiplex identification of viable pathogens in this study (Fig 1). Specifically, an optimized universal RNA extraction and DNA digestion method is developed to simplify and standardize the RNA preparation for both Gram-positive and Gram-negative bacteria. Direct metatranscriptomic RNA sequencing and multiplex RT-PCR amplicon sequencing were evaluated and compared using a cocktail culture of *Escherichia coli* O157:H7 (*E. coli* O157:H7), *Salmonella enteritidis* (*S. enteritidis*), and *Listeria monocytogenes* (*L. monocytogenes*) in both standard general-purpose media and food model. The three bacteria were selected based on their impact on economic loss or prevalence in recent outbreaks.

**Fig 1.**
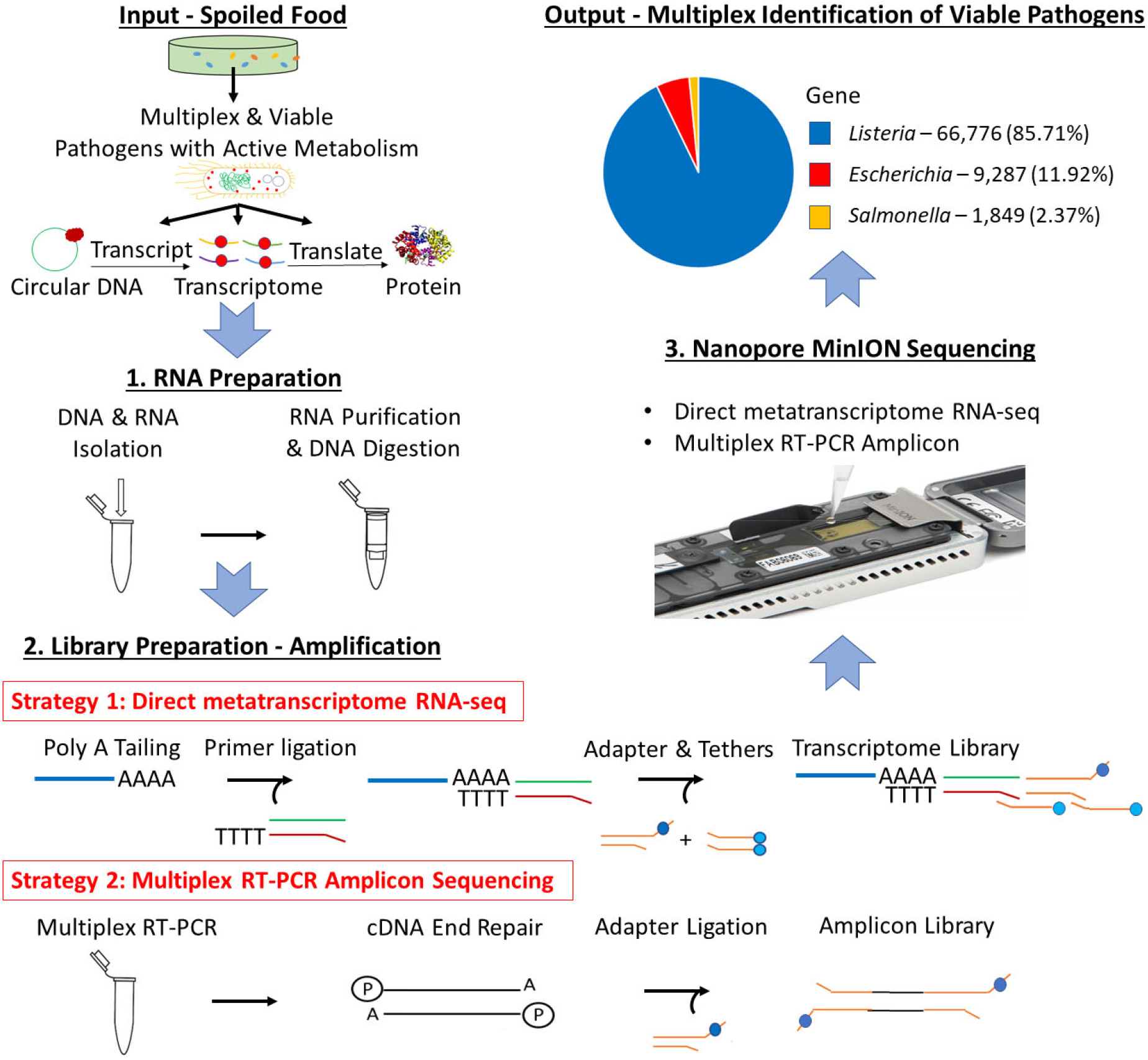
Scheme of multiplex identification of viable pathogens on Nanopore MinION.

## Materials and methods

### Bacterial strains and culturing

*E. coli* O157:H7 (ATCC 43895), *S. enteritidis* (ATCC 13076), and *L. monocytogenes* (ATCC 19115) were acquired from ATCC (Manassas, VA). The three bacteria were cultured using Brain Heart Infusion (BHI) broth and agar (BD, Franklin Lakes, NJ) at 37°C for 24 hours either in separate individual cultures or in cocktail cultures. Romaine lettuce (*Lactuca sativa* L. var. *longifolia*) juice extract (LJE) was used as a food model in this study. Romaine LJE was prepared according to our previous publications (35). Briefly, 250g fresh Romaine lettuce heart (Fresh Express) and 200ml DI Water was blended in a Waring 7011G Commercial Blender for 1 minute. The blended mixture was then filtered through Büchner funnel with P5 filter paper. The filtrate was centrifuged at 4500 rpm for 10 minutes (low speed centrifugation), the supernatant from low speed centrifugation was then centrifuged at 6500 rpm for 30 min (high speed centrifugation). High speed centrifugation supernatant was then filtered through 0.2 micro filter membrane (vacuum filter 0.2-micron, Fisher Scientific) and diluted to 4% using sterilized DI water (COD=800 ppm) to grow bacteria.

Cocktail culture of *E. coli* O157:H7, *S. enteritidis*, and *L. monocytogenes* in Brain Heart Infusion (BHI) or LJE were obtained by inoculating appropriate volume of 24-hour stock culture of the individual bacteria to achieve the initial concentration shown in S2 Table. The concentration for each bacteria was determined by plate counting methods using selective agars. Oxford *Listeria* selective agar base (Oxford formulation) with Oxford modified *Listeria* selective supplement was used for the selective quantification of *L. monocytogenes*. MacConkey agar (BD, Franklin Lakes, NJ) was used to differentiate and quantify *E. coli* O157:H7 and *S. enteritidis*. Both cultures were incubated at 37°C for 24 hours before quantification using an automated plate counter (Scan 300, Interscience Laboratories Inc., Woburn, MA).

### Verification of mRNA as biomarkers for bacteria viability

*E. coli* O157:H7 was selected as a model organism to demonstrate that mRNA is a valid indicator for bacteria viability. Aliquot of overnight *E. coli* O157:H7 culture was inoculated at 3-log CFU/mL in BHI broth and incubated at 37 °C. Culture was sampled at 0, 4, 8, 24 and 72 hours, fractions from the sample culture were taken and plated on BHI agar to determine viable bacteria counts. Remaining fractions were used for DNA and RNA purification using DNeasy blood and tissue kit (Qiagen, Germantown, MD) and Monarch Total RNA Miniprep Kit (New England Biolabs, Ipswich, MA) following the supplier protocols, respectively. In RNA preparation, to lyse the cells, the cell pellet obtained from initial centrifugation was incubated at 37°C for 1 hour with 300 rpm mixing in 250 μL 3 mg/mL lysozyme (Alfa Aesar, Haverhill, MA) in Tris-EDTA buffer (Sigma-Aldrich, St. Louis, MO). Purified DNA and RNA from these four time points were quantified using Qubit dsDNA HS Assay Kit and Qubit RNA HS Assay Kit (Invitrogen, Carlsbad, CA), and also quantified using NEB Luna Universal qPCR Master Mix and Luna Universal One-Step RT-qPCR Kit (New England Biolabs, Ipswich, MA) for qPCR and RT-qPCR test, following supplier protocols, with Biorad CFX-96 Touch real time PCR detection system. The primer pairs used in this test were designated as Stx1A and sequence was listed in S3 Table. *E. coli* O157:H7 inactivated with 13.4 mmol/L of sodium hypochlorite was used as the negative control to test whether mRNA and/or DNA can be used as viability biomarkers (36).

### Direct metatranscriptome RNA-seq on Nanopore MinION

One dimensional direct metatranscriptome RNA-seq was performed using 24-hour cocktail culture of *E. coli* O157:H7, *S. enteritidis*, and *L. monocytogenes* in BHI or LJE. RNA was extracted from the cocktail culture by Monarch Total RNA Miniprep Kit including DNase I (NEB, Ipswich, MA) which was confirmed by multiplex RT-PCR and gel electrophoresis. DNA was completed digested using DNase I (NEB # T2010S, working concentration: 0.1U/μl), which was verified by multiplex PCR and gel electrophoresis. The primers stx, invA and LisA2 were selected for *E. coli* O157:H7, *S. enteritidis*, and *L. monocytogenes,* respectively. RT-PCR products were analyzed using gel electrophoresis with 1.2% agarose gel. NEB One Taq One-Step RT-PCR Kit (NEB # E5310S) was used for nucleic acid amplification. The thermal cycler condition was: 1. Reverse transcription at 48°C for 15 minutes; 2. Initial denaturation at 94°C for 1 minute; 3. Denaturation at 94°C for 15 sec, annealing at 53°C for 30 sec, extension at 68°C for 40 sec with 40 cycles; 4. Final extension at 68°C for 5 minutes.

The prepared RNA samples were further modified with poly(A) tailing and library preparation by following suppliers’ protocols. Direct metatranscriptome RNA-seq was developed based on supplier’s direct RNA-seq protocol (RNA Kit SQK-RNA001, Nanopore, Oxford, United Kingdom). The MinION flow cell was primed using a priming mix, and then 75 μl of sample was loaded to the SpotON sample port dropwise to avoid bubbles. After adding the sample, MinKNOW software was initiated to start a sequencing run. Cocktail culture inactivated with 13.4 mmol/L of sodium hypochlorite was used as the negative control to test whether metatranscriptome sequencing can eliminate false positive identification.

### Multiplex RT-PCR amplicon sequencing on Nanopore MinION

Multiplex RT-PCR amplicon sequencing was conducted using 4-hour cocktail culture of *E. coli* O157:H7, *S. enteritidis*, and *L. monocytogenes* in BHI or LJE. RNA extraction and DNA digestion were performed and verified using the same protocols listed above. Major virulent genes selected in this study include: *stx* and *stx1A* localized to the lambdoid prophages H19B and H19J in *E. coli* O157:H7 (37–39); *invA*, a critical component of the *Salmonella* pathogenicity island 1 (SPI1) in *S. enteritidis* (39, 40); and *inlA,* encoded internalins genes in the *inlAB* operon outside the *Listeria* pathogenicity island 1 (LIPI-1) in *L. monocytogenes* (39, 41, 42).

Prior to RT-PCR amplicon sequencing, an end repair/A tailing step (NEBNext End Repair/dA-tailing Module) was carried out for the RT-PCR products, followed by a ligation step using NEB Ultra II ligation master mix. Oxford Nanopore Ligation Sequencing Kit SQK-LSK108 and Library Loading Bead Kit EXP-LLB001 were used for the library preparation of RT-PCR amplicon. To validate Nanopore DNA sequencing results, the RT-PCR products were also sequenced using an Applied Biosystems 3130xl genetic analyzer by following a protocol at NEB. The DNA sequencing data collected by the sequencer was analyzed using EPI2ME and MG-RAST to confirm the identities of the three bacteria. Cocktail culture inactivated with 13.4 mmol/L of sodium hypochlorite was used as the negative control to test whether multiplex RT-PCR amplicon sequencing can eliminate false positive identification. Detailed protocols for RT-PCR are provided in the supplemental material.

### Data analysis and bioinformatics

Sequencing reads were base-called via the local base-calling algorithm with MinKNOW software (v. 1.4.3). All FASTQ files of passed base-called reads were collected and combined to one file for analysis. EPI2ME, MG-RAST and MEGAN were used for metagenomics and taxonomic analysis.

## Results

### Verification of mRNA as biomarkers for bacteria viability

Fig 2 showed qPCR, RT-qPCR results of *E. coli* O157:H7 samples collected from 5 time points. *E. coli* O157:H7 growth curve (Fig 2A) resembles a typical microbial growth curve with an exponential phase from 0 to 24 hours and a stationary phase from 24 to 72 hours. Bacterial counts at 72 hours showed a slight decrease from 24 hours, which could indicate the start of death phase. The RT-qPCR of mRNA collected at different time points (Fig 2B) showed that the greatest amount of RNA was found in 8-hour and 24-hour samples, followed by a decline of mRNA concentration at 72 hours. A high alignment between mRNA concentration and viable cell density can be established between Fig 2A and 2B. The results indicate mRNA has good correlation with viable bacterial count. The melt curve (Fig 2E) showed 5 peaks from the 5 different time points, which indicate that the same mRNA was amplified. In negative control, no colony was identified on BHI agar and no amplicon was detected by gel electrophoresis.

**Fig 2.**
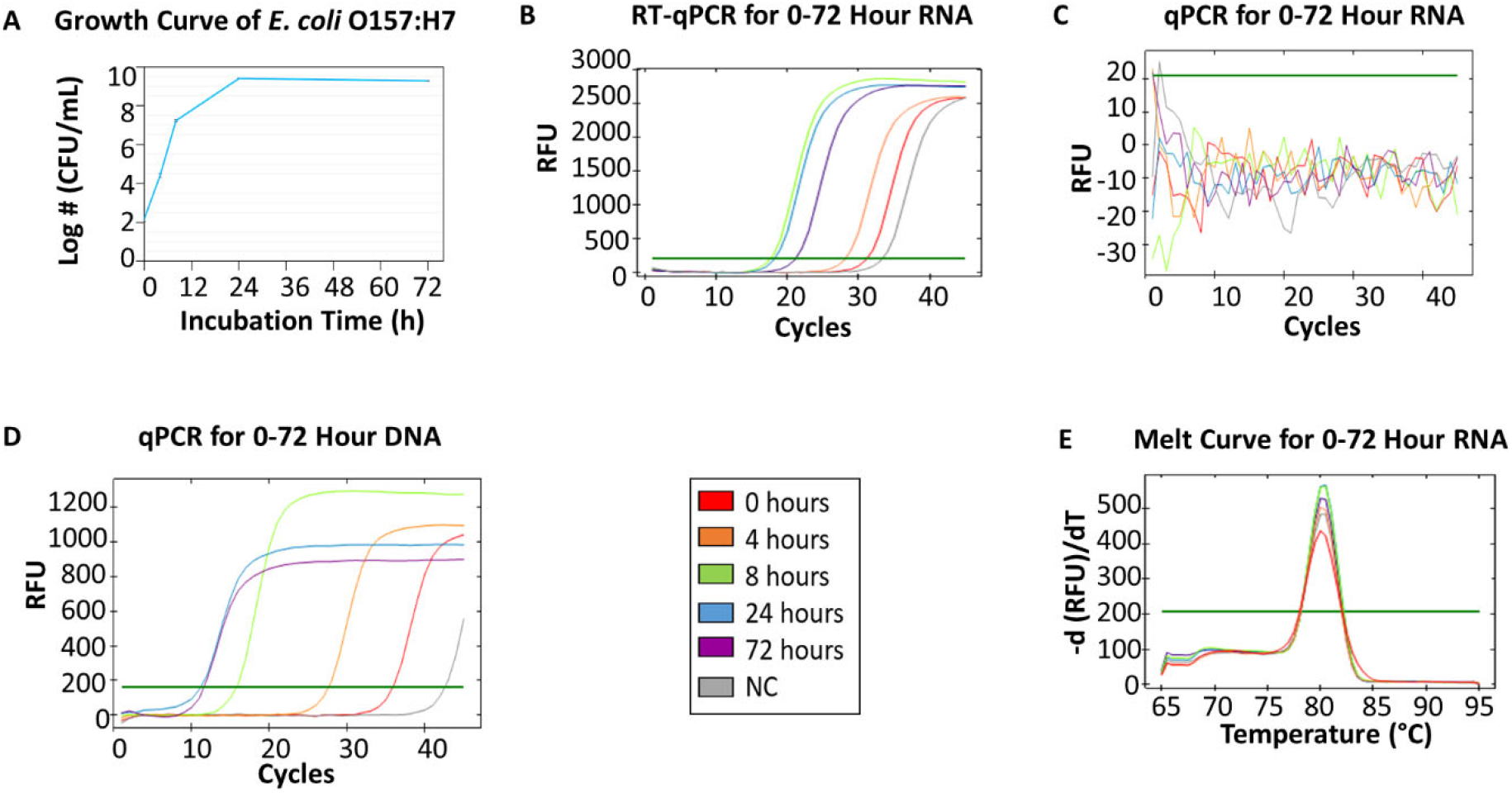
RT-qPCR and qPCR of *E. coli* O157:H7 from 0, 4, 8, 24 and 72 h growth in BHI. (A) The growth curve of *E. coli* O157:H7 at 0, 4, 8, 24, 72 h in BHI. The initial concentration was 3-log CFU/mL. (B) RT-qPCR for RNA collected from 5 time points. (C) qPCR for 0-72 h RNA as the negative control (NC) for DNA contamination – no DNA contamination was found in those samples. (D) qPCR for DNA collected from 5 time points. (E) The melting curve analysis of RT-qPCR for 0-72 h RNA.

The qPCR of *E. coli* O157:H7 DNA (Fig 2D) showed that the amount of DNA in 72-hour samples was greater than the amount in 24-hour samples, which contradicted the data from the viable bacterial counts. This indicated DNA accumulation from nonviable cells was present in 72-hour samples, which was consistent with other studies (43, 44). Hence, DNA was not a great indicator of bacteria viability. Additionally, the same qPCR amplicon was detected by gel electrophoresis in the negative control of sodium hypochlorite treated *E. coli* O157:H7. Therefore, the results demonstrate that the global transcriptome, especially mRNA, of bacteria could be a robust indicator of cell viability.

### Direct metatranscriptome RNA-seq on Nanopore MinION

Table 1 showed the results of direct metatranscriptome RNA-seq of *E. coli* O157:H7, *S. enteritidis* and *L. monocytogenes* cocktail in BHI and LJE 24-hour culture using different bioinformatics pipelines. Both EPI2ME, MG-RAST, and MEGAN miss-identified the three pathogens as other species (Table 2). MEGAN with non-rRNA mapping successfully identified the three bacteria without miss-identification as *Listeria, E. coli* and *Salmonella* at 91.1%, 5.4% and 3.6% in BHI and 67.5%, 20% and 12.5% in LJE, respectively (Table 2). The results agreed with plate counting confirmation that all three bacteria were present (S2 Table). The mean read-length was close to 1,200 bp (Table 1 and S2 Fig.), which agrees with the size of 16S RNA in bacteria. The average quality scores were 7.8 in BHI and 7.9 in LJE. There was no miss-identification of the bacteria using a quality score cut-off at 7.0 using MEGAN analysis with non-rRNA mapping. No false positive identification of any bacteria was found in the negative control of sodium hypochlorite treated cocktail culture (negative control). Therefore, the results strongly support that direct metatranscriptome RNA-seq on Nanopore MinION can achieve multiplex identification of viable pathogens.

**Table 1.**
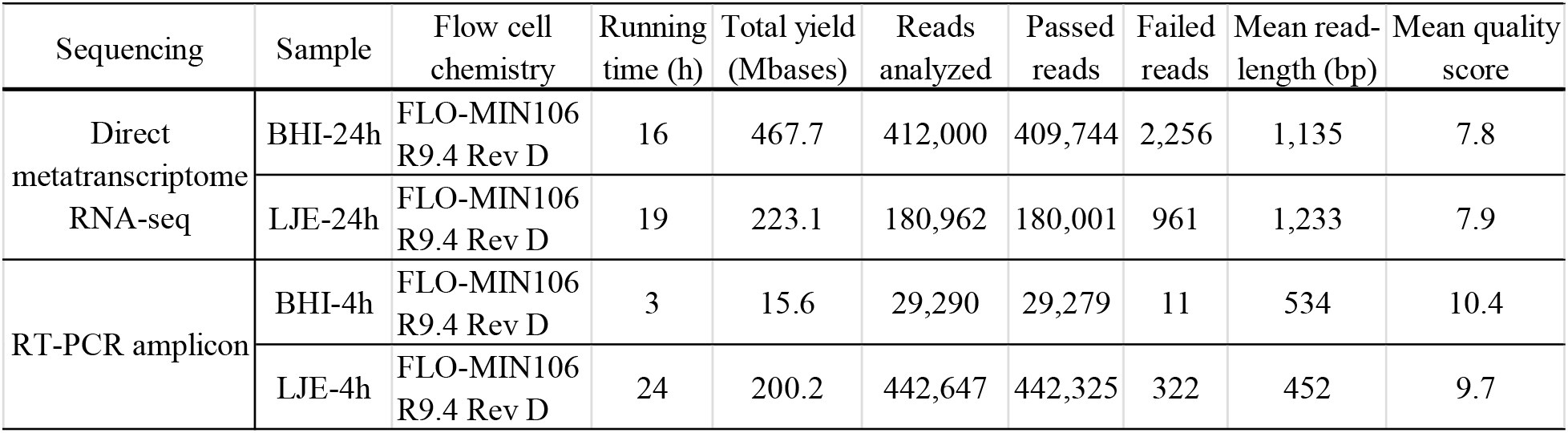
Direct metatranscriptome RNA-seq and amplicon sequencing of cocktail bacterial culture on Nanopore platform.

| Sequencing | Sample | Flow cell chemistry | Running time (h) | Total yield (Mbases) | Reads analyzed | Passed reads | Failed reads | Mean read-length (bp) | Mean quality score |
| --- | --- | --- | --- | --- | --- | --- | --- | --- | --- |
| Direct metatranscriptome RNA-seq | BHI-24h | FLO-MIN106 R9.4 Rev D | 16 | 467.7 | 412,000 | 409,744 | 2,256 | 1,135 | 7.8 |
|  | LJE-24h | FLO-MIN106 R9.4 Rev D | 19 | 223.1 | 180,962 | 180,001 | 961 | 1,233 | 7.9 |
| RT-PCR amplicon | BHI-4h | FLO-MIN106 R9.4 Rev D | 3 | 15.6 | 29,290 | 29,279 | 11 | 534 | 10.4 |
|  | LJE-4h | FLO-MIN106 R9.4 Rev D | 24 | 200.2 | 442,647 | 442,325 | 322 | 452 | 9.7 |

**Table 2.** Results of MinION R9.4 Rev D direct metatranscriptome RNA-seq and RT-PCR amplicon sequencing for BHI and LJE samples collected from 4-hour and 24-hour culture with different initial growth concentration via different bioinformatic pipelines.*

| Sequencing | Sample | Bioinformatic pipeline |  |  |  |  |  |  |  |
| --- | --- | --- | --- | --- | --- | --- | --- | --- | --- |
|  |  | MinKNOW/EPI2ME |  | MG-RAST |  | MEGAN |  | MEGAN-rRNA excluded |  |
|  |  | Taxon | Cumulative reads | Taxon | Cumulative reads | Taxon | Percentage | Taxon | Percentage |
| Direct metatranscriptome RNA-seq | BHI-24h | *Bacillus* | 65,746 (37.6%) | *Listeria* | 75,562 (85.28%) | *Bacillus* | 63.6% | ***Listeria*** | **91.1%** |
|  |  | *Listeria* | 54140 (31.0%) | *Escherichia* | 4,560 (5.15%) | *Listeria* | 29.5% | ***Escherichia*** | **5.4%** |
|  |  | *Escherichia* | 29312 (16.8%) | *Salmonella* | 1,281 (1.45%) | *Escherichia* | 4.5% | ***Salmonella*** | **3.6%** |
|  |  | *Lactobacillus* | 10718 (6.1%) | *Coptotermes* | 1,237 (1.40%) | *Salmonella* | 2.3% | - | - |
|  |  | *Staphylococcus* | 8772 (5.0%) | *Bacillus* | 952 (1.07%) | - | - | - | - |
|  |  | *Salmonella* | 6155 (3.5%) | *Tetragenococcus* | 435 (0.49%) | - | - | - | - |
|  | LJE-24h | *Bacillus* | 25434 (34.5%) | *Listeria* | 66,776 (76.55%) | *Bacillus* | 63.6% | ***Listeria*** | **67.5%** |
|  |  | *Escherichia* | 23084 (31.3%) | *Escherichia* | 9,287 (10.65%) | *Listeria* | 29.5% | ***Escherichia*** | **20.0%** |
|  |  | *Listeria* | 16751 (22.7%) | *Salmonella* | 1,849 (2.12%) | *Escherichia* | 4.5% | ***Salmonella*** | **12.5%** |
|  |  | *Salmonella* | 4943 (6.7%) | *Bacillus* | 1,028 (1.18%) | *Salmonella* | 2.3% | - | - |
|  |  | *Edwardsiella* | 3515 (4.8%) | *Lactobacillus* | 728 (0.83%) | - | - | - | - |
| RT-PCR amplicon | BHI-4h | ***Escherichia*** | **5770 (84.7%)** | *Escherichia* | 1,342 (70.19%) | - | - | - | - |
|  |  | ***Salmonella*** | **931 (13.7%)** | *Salmonella* | 474 (24.79%) | - | - | - | - |
|  |  | *Listeria* | 113 (1.7%) | *Enterobacter* | 43 (2.25%) | - | - | - | - |
|  |  | - | - | *Listeria* | 20 (1.05%) | - | - | - | - |
|  | LJE-4h | ***Escherichia*** | **77437 (50.9%)** | *Escherichia* | 1,852 (88.11%) | - | - | - | - |
|  |  | ***Salmonella*** | **66098 (43.4%)** | *Listeria* | 179 (8.52%) | - | - | - | - |
|  |  | *Listeria* | 8711 (5.7%) | *Enterobacter* | 45 (2.14%) | - | - | - | - |

\* Bold indicates the optimal bioinformatics pipeline without miss-identification.

### Multiplex RT-PCR amplicon sequencing on Nanopore MinION

Similarly, multiplex RT-PCR amplicon sequencing also successfully identified the three bacteria in the 4-hour cocktail culture sample (Fig 3). *E. coli* O157:H7, *S. enteritidis* and *L. monocytogenes* were observed in a real-time phylogenetic tree generated by EPI2ME in less than 15 minutes and the distribution was respectively 84.7%, 13.7% and 1.7% in BHI sample, and 50.9%, 43.4% and 5.7% in LJE sample (Fig 3A, 4B and Table 2). The average read quality was 10.4 and 9.7. A total of 29,279 reads were analyzed in BHI culture with a 3-hour running time (early termination due to high quality score) and 442,325 reads in LJE cocktail culture with a 24-hour running time (Table 1). The average sequence length was 534 bp in BHI sample and 432 bp in LJE sample (S3 Table and S2 Fig.) which was consistent with multiplex RT-PCR products (520, 244 and 153 bp) (39–42). No false positive identification of any bacteria was found in the negative control of sodium hypochlorite treated cocktail culture.

**Fig 3.**
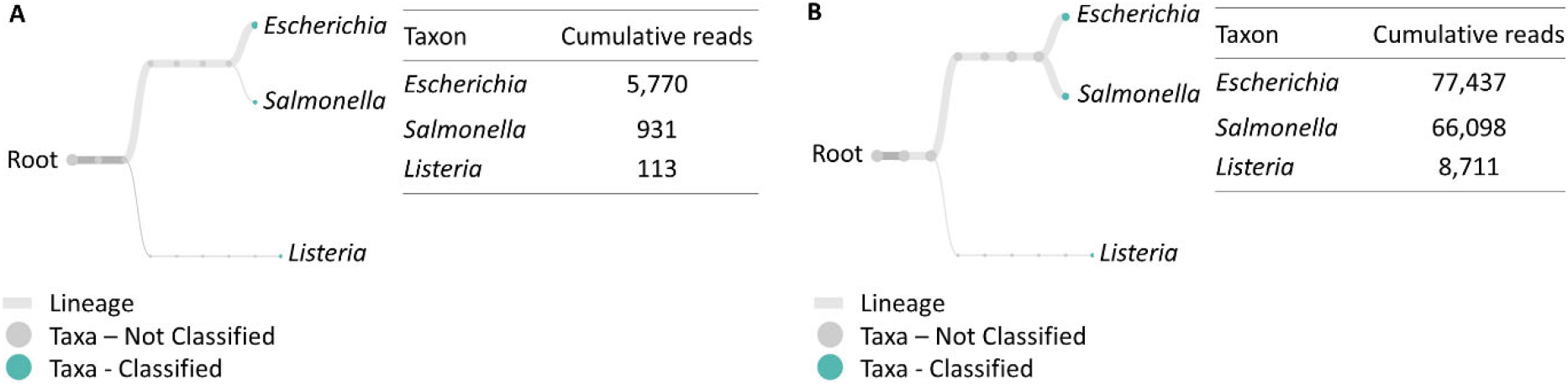
Taxonomic and genus level bacterial classification of MinION R9.4 Rev D multiplex RT-PCR amplicon sequencing. (A) Taxonomy tree of BHI 334 4-hour sample generated by EPI2ME. (B) Taxonomy tree of LJE 334 4-hour sample generated by EPI2ME.

### Quality control and comparison of bioinformatic pipelines

In this study, the quality score of direct metatranscriptome RNA-seq was 7.8 and 7.9, while 10.4 and 9.7 for multiplex RT-PCR amplicon sequencing from MinKNOW QC report (Table 1). Raw data was collected by nanopore real-time sequencing software MinKNOW and analyzed with different bioinformatic databases and pipelines. In metatranscriptomic direct RNA-seq, MinKNOW miss-identified *Bacillus* as the top genus in both the BHI and LJE samples (Table 2). This error could be caused by the similarity between *Listeria* and *Bacillus*, especially with their housekeeping genes and rRNA (45, 46). Although MG-RAST eliminated this misreading, other untargeted bacteria counted for a close proportion to *S. enteritidis* (1.5%) (Table 2). MEGAN was able to eliminate other untargeted bacteria except *Bacillus* (still misidentified as 63.6%), which again was likely caused by rRNA or other housekeeping genes. Therefore, MEGAN with non-rRNA mapping was performed and successfully identified all three primary bacteria of *Listeria*, *E. coli* O157:H7 and *Salmonella* without any miss-identification (Table 2). The results of multiplex RT-PCR sequencing showed that three targeted bacteria were anchored accurately by MinKNOW (Table 2), and no miss-identification appeared in the results.

### Gel electrophoresis of RT-PCR and PCR amplicon

RT-PCR was used to verify the presence of all three bacteria in the cocktail culture, and PCR was used to verify the complete removal of DNA contamination using the protocol described above.

S3A Fig. shows the verification of 24-hour LJE cocktail culture, which was used in metatranscriptomic direct RNA-seq. The results showed the RT-PCR product of three expected bands for *stx*, *invA* and *inlA* in the 24-hour cocktail culture, which confirms the presence of all three target pathogens. No bands appeared on negative controls using only PCR without the RT step, which indicates the absence of DNA contamination. Further validation was performed for the 4-hour LJE cocktail culture sample, which was used in the multiplex RT-PCR amplicon sequencing. S3B Fig. shows multiplex RT-PCR amplicon with three bands. The sizes of the amplicons are consistent with previous reports of 520 (*stx*), 244 (*invA*) and 153 (*inlA*) bp (S3B Fig. line 2). No RT-PCR products were detected in the negative control (line 3, 4 and 5) using only PCR without the RT step, which indicates that there was no DNA contamination in the sample.

### Discussion Viability and multiplex identification

In this study, RNA-enabled Nanopore sequencing is evaluated, for the first time, for its potential in achieving multiplex identification of viable pathogens. The optimized universal RNA extraction and DNA digestion method was developed to simplify and standardize the RNA preparation for both Gram-positive and Gram-negative bacteria. Direct metatranscriptome RNA-seq and multiplex RT-PCR amplicon sequencing were evaluated and compared using a cocktail culture of *E. coli* O157:H7, *S. enteritidis*, and *L. monocytogenes* in both standard general-purpose media and food model. False positives are a major issue for DNA-based approaches. This is due to the inability to differentiate DNA molecules in viable bacterial cells from the genomic background, which is comprised of stable DNA molecules from the microbiota, the food matrices, and dead pathogens inactivated during food processing and storage. Both approaches developed in this study only utilize RNA, especially mRNA, as the ultimate sequencing target, which eliminated false positive identification typically caused by DNA contamination. Random and unknown threats from multiple infectious bacteria poses significant threat to the safety and security of food supply worldwide. A feasible strategy should enable multiplex identification without the need to customize for an individual threat. Therefore, the developed universal protocol is applicable to both Gram-positive and Gram-negative bacteria. RNA from multiple pathogens in one food sample can be collected from one extraction and library preparation step, followed by the universal sequencing protocol.

### Comparison of direct metatranscriptome RNA-seq and multiplex RT-PCR amplicon sequencing

The developed method successfully identified all three bacteria from cocktail culture in BHI and LJE by direct metatranscriptome RNA-seq and multiplex RT-PCR amplicon sequencing. Nonetheless, the two sequencing approaches entail different capacities and challenges. Direct metatranscriptome RNA-seq does not require assay customization for an individual biohazard if the bioinformatic database includes the target microbiota. Multiplex RT-PCR amplicons comprise the target gene copies, and they can be easily captured by the motor membrane protein when passing through the nanopores. As a result, it shows higher accuracy, greater quality score, better quality control, and less turnaround time. The two strategies result in different read length. In this study, we extracted total bacterial RNA that is comprised of a majority of rRNA and a small number of mRNA and tRNA for nanopore sequencing. The bioanalyzer results showed that a majority of RNA from *E. coli* O157:H7, *S. enteritidis* and *L. monocytogenes* cocktail culture were 16S RNA with 1250 – 2100 nt, and 23S RNA with 2250 – 3950 nt, respectively. Direct metatranscriptome RNA-seq sequencing successfully identified all three bacteria. The RNA read length ranged from 0 to 3000 nt, with the most abundant read length between 400-1600 nt. The read length from direct metatranscriptome RNA-seq is approximately the full length of RNA (47, 48). Thus, this method can provide approximal full-length RNA sequence. However, in multiplex RT-PCR amplicon sequencing, the amplicons for each bacteria have different expected sizes, which were observed in the sequencing read length results. In the real-time analysis of multiplex RT-PCR amplicon sequencing, all three bacteria were identified within 15 minutes and the resulting read lengths were 510 bp (*stx*), 244 bp (*invA*) and 153 bp (*inlA*), respectively. Some reports suggest that Nanopore excels in long RNA reads up to thousands of nt, and sequencing of short reads tends to be more challenging due to their higher and non-uniform error profiles, which might result in a large fraction of reads remaining unmapped or unused (49–51). However, amplicon sequencing showed less error than metatranscriptomic direct RNA-seq. Multiplex RT-PCR amplicon sequencing successfully identified all three target bacteria using MinKNOW (Table 2) in real time. Direct metatranscriptome RNA-seq experienced miss-identification if selecting the wrong bioinformatic databases or pipelines. Both methods are comparable in their total turnaround time. Direct metatranscriptome RNA-seq does not include an additional RT-PCR step, but the library preparation, bioinformatic analysis, and mapping could easily offset the time difference. The total turnaround time for direct metatranscriptome RNA-seq is approximately 6.5 hours, which includes RNA purification (3.5 hours), library preparation (1.5 hours), Nanopore sequencing (1 hour), and bioinformatic analysis (0.5 hour). Multiplex amplicon sequencing takes approximately 6 hours, which includes RNA purification (3.5 hours), RT-PCR (2 hours), library preparation (0.5 hour), Nanopore sequencing (15 minutes), and bioinformatics analysis (0.5 hour).

Multiplex RT-PCR amplicon sequencing requires substantially less RNA input, which could translate into less microbial input. The method only requires 36.5 ng RNA input for multiplex RT-PCR, and 33.8 ng amplicon for library preparation and sequencing (S2 Table). The amplicon sequencing method is more sensitive and could be applicable for food commodities with low bacterial loading around 10^1^-10^4^ CFU/g. 500 ng RNA input on Nanopore MinION is recommended by the supplier for metatranscriptomic direct RNA-seq. However, significant RNA loss was observed during the library preparation due to the three purification steps. The initial purified RNA concentration before library preparation was 3490 ng and 1338 ng in BHI and LJE respectively, and only 744 ng and 130 ng were yielded for Nanopore sequencing. RNA loss can be as high as 80-90%, which significantly restricted sensitivity of the assay. Redesign of the library preparation protocol to minimize RNA loss can have profound significance for assay sensitivity and feasibility for clinical applications. The two strategies pose different levels of complexities. Direct metatranscriptome RNA-seq may be applicable in foods with a complex microbiome (e.g., cultured food). Direct metatranscriptome RNA-seq does not require assay customization for an individual biohazard, if the bioinformatic database includes the target microbiota. The multiplex RT-PCR amplicon sequencing requires complex primer design and validation. Not all RT-PCR primers work in multiplex RT-PCR, due to potential primer interaction, nonspecific amplification, and amplification bias. The amplicon sequencing may be more suitable for high-throughput and continuous monitoring of foodborne pathogens with high risk factors.

### Comparison between different bioinformatic pipelines

Bioinformatic analysis has significant impact to the accuracy of Nanopore sequencing. Different computational pipelines of the same nanopore data may lead to different results. Normally, MinION pipeline contains primer trimming, alignment, variant calling and consensus generation (52–55), and EPI2ME conducts real-time surveillance of nanopore sequencing. First, reads containing raw data are base called by MinKNOW, and then extracted into a FASTQ file for mapping to reference transcriptome or genome (56–58), aligned to sequence via primer trimming and coverage normalization. During this process, low quality or low coverage reads (read hit) are filtered out to generate final sequence for BLAST in NCBI. MinION chemistry provides a simplified and rapid report of nanopore running, including read number, read length, cumulative read, taxonomy tree and quality control. MEGAN and MG-RAST are popular software or service for metagenome or metatranscriptome analysis. The similarity between them is that they perform computational analysis of multiple datasets for taxonomic content based on family and gene level. In contrast, MEGAN is able to perform taxonomical, functional and interactive analyses, which is the comparison of taxonomic and functional contents based on the SEED hierarchy and KEGG pathways (59, 60). In this study, MG-RAST was used in taxonomic analysis, and MEGAN was selected for mRNA analysis in addition to MinION. Both show a higher accuracy compared with MinION for metatranscriptome sequencing. Multiplex RT-PCR amplicon obtained a rapid and accurate taxonomic content because this nanopore sequencing method poses a high sensitivity. In addition, adequate and complete BLAST database may further improve the accuracy, rapidness, and quality for the multiplex identification of viable pathogens in food. In conclusion, novel strategies are developed through direct metatranscriptome RNA-seq and multiplex RT-PCR amplicon sequencing on Nanopore MinION to achieve real-time multiplex identification of viable pathogens in food. This study reports an optimized universal Nanopore sample extraction and library preparation protocol applicable to both Gram-positive and Gram-negative. Further evaluation and validation confirmed the accuracy of direct metatranscriptome RNA-seq and multiplex RT-PCR amplicon sequencing using Sanger sequencing and selective media. The study also included a comparison of different bioinformatic pipelines for metatranscriptomic and amplicon genomic analysis. In addition, direct metatranscriptome RNA-seq and RT-PCR amplicon sequencing were compared for their respective advantages in sample inputs, accuracy, sensitivity, and time effectiveness for potential applications. Both direct metatranscriptome RNA-seq and multiplex RT-PCR amplicon sequencing need more development to address some pressing challenges. A) Optimization of direct metatranscriptome RNA-seq sequencing may include minimizing RNA loss in the library preparation step; comparison of bioinformatic pipelines to eliminate miss-identified and unclassified targets; cross-domain identification of prokaryotes, eukaryotes, and viruses. B) Multiplex RT-PCR amplicon sequencing can benefit from: multiplex primer development; inclusivity/exclusivity evaluations; reduced amplification bias. To the best of our knowledge, this is the first report of metatranscriptome sequencing of cocktail microbial RNAs on the emerging Nanopore platform. Direct RNA-seq and RT-PCR amplicons sequencing of metatranscriptome enable the direct identification of nucleotide analogs in RNAs, which is highly informative for determining microbial identities while detecting ecologically relevant processes. The information pertained in this study could be important for future revelatory research including predicting antibiotic resistance, elucidating host-pathogen interaction, prognosing disease progression, and investigating microbial ecology, etc.

## Acknowledgments

The project is partially supported by the U.S. Department of Agriculture (S51600000035794). Special appreciation goes to LiTing Chiu and Tong Wu for their support in this project.

