## Supplemental Materials for "Direct Metatranscriptome RNA-seq and Multiplex RT-PCR Amplicon Sequencing on Nanopore MinION – Promising Strategies for Multiplex Identification of Viable Pathogens in Food"

TABLE S1 Summary of conventional cultivation and commercial rapid detection methods.

| Method | Rapidness (hours)* | VBNC* | Portability* | Viability* | Multiplex w/o customization* |
|---|---|---|---|---|---|
| Most probable number | **24–48** | **No** | Yes | Yes | Yes |
| Viable counts (e.g., Petrifilm™) | **24–72** | **No** | Yes | Yes | Yes |
| Lateral flow | <1 | **No** | Yes | **No** | **No** |
| DEFT/SPC | <1 | **No** | **No** | **No** | **No** |
| Immunoassay | 1–4 | **Enrichment required** | Yes | **Enrichment required** | **No** |
| Flow cytometry | <1 | Yes | **No** | Yes | **No** |
| MALDI-TOF | 1 | **No** | **No** | Yes | Yes |
| DNA gene / PCR | ~4 | Yes / **False positive** | Yes | **No / False positive** | **No** |
| RNA transcript / RT-PCR | ~4 | Yes / **False positive** | Yes | Yes / **False positive** | **No** |
| Next generation sequencing (NGS) | 1–4 | Yes | **No** | Yes | Yes |
| **Nanopore** | **1–4** | **Yes** | **Yes** | **Yes** | **Yes** |

* Bold indicates major limitations.

TABLE S2 Nanopore sequencing input amount of RNA and RT-PCR amplicon (cDNA). CFU number was measured by plating count and nucleotide concentration was tested by Qubit assay. *Ec*, *Se* and *Lm* indicated *E. coli* O157:H7, *S. enteritidis* and *L. monocytogenes*, respectively. The bacteria culture condition was 37°C incubation.

| Sequencing | Direct metatranscriptome RNA-seq | | | | | | RT-PCR amplicon | | | | | |
|---|---|---|---|---|---|---|---|---|---|---|---|---|
| Sample | BHI 336 24h | | | LJE 336 24h | | | BHI 334 4h | | | LJE 334 4h | | |
| Initial culture conc. (Log CFU/mL) | *Ec* | *Se* | *Lm* | *Ec* | *Se* | *Lm* | *Ec* | *Se* | *Lm* | *Ec* | *Se* | *Lm* |
| | 2.9 | 3.1 | 4.3 | 3.5 | 3.5 | 5.9 | 3.6 | 3.5 | 4.7 | 3.6 | 3.3 | 5.5 |
| 24h/4h culture conc. (Log CFU/mL) | *Ec* | *Se* | *Lm* | *Ec* | *Se* | *Lm* | *Ec* | *Se* | *Lm* | *Ec* | *Se* | *Lm* |
| | 8.7 | 9.2 | 7.3 | 8.6 | 8.0 | 5.4 | 5.3 | 5.0 | 5.7 | 4.5 | 3.6 | 4.5 |
| Sample size (mL) | 5.3 | | | 5.0 | | | 0.2 | | | 0.2 | | |
| RNA input for multiplex RT-PCR (ng) | - | | | - | | | 72.0 | | | 36.5 | | |
| Yield of cDNA from multiplex RT-PCR (ng) | - | | | - | | | 1267.5 | | | 705.2 | | |
| RNA/DNA input for library prep (ng) | 3490.2 | | | 1338.0 | | | 33.8 | | | 165.0 | | |
| Yield after Poly A tailing and purification (ng) | 1824.0 | | | 617.5 | | | - | | | - | | |
| Yield after library prep and Nanopore sequencing input (ng) | 744.0 | | | 129.2 | | | <30.0 Too low to detect | | | 125.0 | | |

TABLE S3 Primer information for qPCR, RT-qPCR and multiplex RT-PCR.

| Species | Target Gene* | Primer (5'-3') | Product Size (bp) | Reference |
|---|---|---|---|---|
| *Ec* | *stx* | F:GAGCGAAATAATTTATATGTG<br>R:TGATGATGGCAATTCAGTAT | 520 | (1-3) |
| | *stx1A* | F: TGACAGGATTTGTTAACAGGAC<br>R: TCTGTATTTGCCGAAAACGT | 294 | (1-3) |
| *Se* | *invA* | F:ACAGTGCTCGTTTACGACCTGAAT<br>R:AGACGACTGGTACTGATCGATAAT | 244 | (3, 4) |
| *Lm* | *inlA* | F:GATTAACACGAGTAACGG | 153 | (3, 5, 6) |

* *stx1A* was used in qPCR and RT-qPCR of *E. coli* O157:H7. *stx*, *invA* and *inlA* were used in multiplex RT-PCR of 4-hour BHI and 4-hour LJE cocktail cultures.

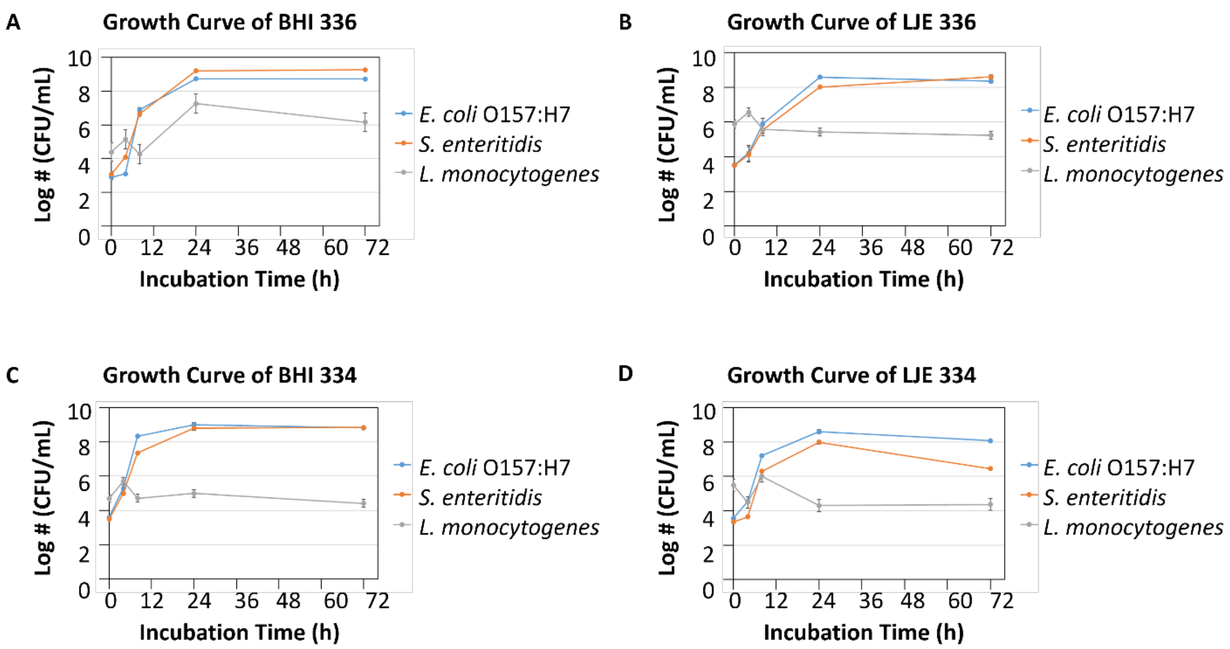


S1 Fig. The growth curve of cocktail culture of *E. coli* O157:H7, *S. enteritidis* and *L. monocytogenes* at 0, 4, 8, 24 and 72 h in BHI and LJE at 37 ºC. (A) The growth curve of cocktail culture of *E. coli* O157:H7 (3-log), *S. enteritidis* (3-log) and *L. monocytogenes* (6-log) at 0, 4, 8, 24 and 72 h in BHI. (B) The growth curve of cocktail culture of *E. coli* O157:H7 (3-log), *S. enteritidis* (3-log) and *L. monocytogenes* (6-log) at 0, 4, 8, 24 and 72 h in LJE. (C) The growth curve of cocktail culture of *E. coli* O157:H7 (3-log), *S. enteritidis* (3-log) and *L. monocytogenes* (4-log) at 0, 4, 8, 24 and 72 h in BHI. (D) The growth curve of cocktail culture of *E. coli* O157:H7 (3-log), *S. enteritidis* (3-log) and *L. monocytogenes* (4-log) at 0, 4, 8, 24 and 72 h in LJE.

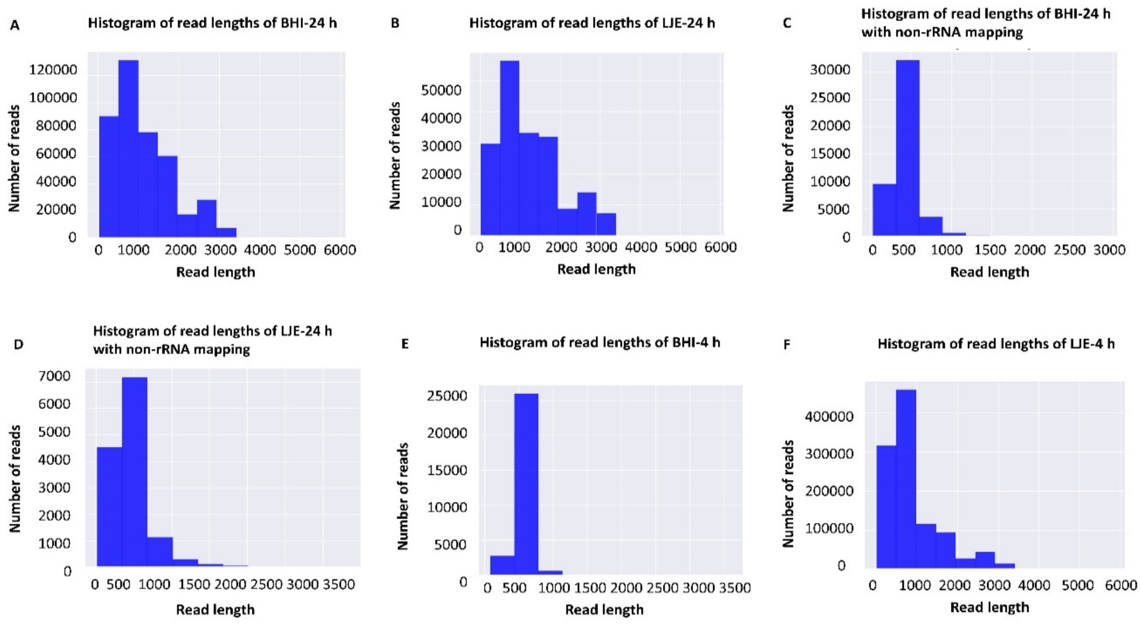


S2 Fig. The number of reads and read lengths of MinION R9.4 Rev D direct metatranscriptome RNA-seq and RT-PCR amplicon sequencing for BHI and LJE samples collected from 4 h and 24 h. Results of direct metatranscriptome RNA-seq. (A) BHI-24 h; (B) LJE-24 h; (C) BHI-24 h with non-rRNA mapping; (D) LJE-24 h with non-rRNA mapping. Results of RT-PCR amplicon sequencing of (E) BHI-4 h; (F) LJE-4 h.

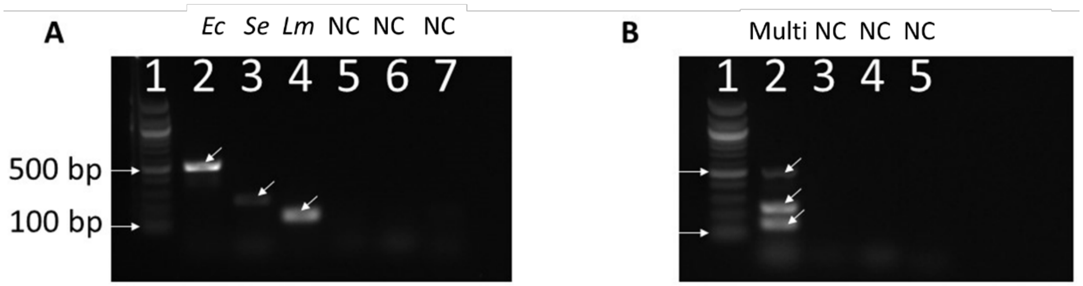


S3 Fig. Gel image of RT-PCR amplicons were used to verify the present of all three bacteria in the cocktail culture, and PCR was used to verify the complete removal of DNA contaminations using the protocol described above. (A) Non-multiplex RT-PCR of *E. coli* O157:H7 (*Ec*), *S. enteritidis* (*Se*) or *L. monocytogenes* (*Lm*) from 24-hour cocktail culture in LJE. From left to right: 1. 100-bp DNA Ladder (NEB); 2. *E. coli* O157:H7 in LJE 24-hour cocktail culture; 3. *S. enteritidis* in LJE 24-hour cocktail culture; 4. *L. monocytogenes* in LJE 24-hour cocktail culture. 5. Negative control (NC) of *E. coli* O157:H7 in LJE 24-hour cocktail culture; 6. Negative control of *S. enteritidis* in LJE 24-hour cocktail culture; 7. Negative control of *L. monocytogenes* in LJE 24-hour cocktail culture. (B) Multiplex RT-PCR (Multi) of *E. coli* O157:H7, *S. enteritidis* and *L. monocytogenes* 4-hour cocktail in LJE. From left to right: 1. 100-bp DNA Ladder; 2. Multiplex RT-PCR of *E. coli* O157:H7, *S. enteritidis* and *L. monocytogenes*; 3. Negative control of *E. coli* O157:H7 (PCR without RT); 4. Negative control of *S. enteritidis* (PCR without RT); 5. Negative control of *L. monocytogenes* (PCR without RT).
